# All-Atom Simulation of the HET-s Prion Replication

**DOI:** 10.1101/2020.05.01.072017

**Authors:** Luca Terruzzi, Giovanni Spagnolli, Alberto Boldrini, Jesús R. Requena, Emiliano Biasini, Pietro Faccioli

## Abstract

Prions are self-replicative protein particles lacking nucleic acids. Originally discovered for causing infectious neurodegenerative disorders, they have also been found to play several physiological roles in a variety of species. Functional and pathogenic prions share a common mechanism of replication, characterized by the ability of an amyloid conformer to propagate by inducing the conversion of its physiological, soluble counterpart. In this work, we focus on the propagation of the prion forming domain of HET-s, a physiological fungal prion for which high-resolution structural data are available. Since time-resolved biophysical experiments cannot yield a full reconstruction of prion replication, we resort to computational methods. To overcome the computational limitations of plain Molecular Dynamics (MD) simulations, we adopt a special type of biased dynamics called ratchet-and-pawl MD (rMD). The accuracy of this enhanced path sampling protocol strongly depends on the choice of the collective variable (CV) used to define the biasing force. Since for prion propagation a reliable reaction coordinate (RC) is not yet available, we resort to the recently developed Self-Consistent Path Sampling (SCPS). Indeed, in such an approach the CV where the biasing force is applied is not heuristically postulated but is calculated through an iterative refinement procedure. Our atomistic reconstruction of the HET-s replication shows remarkable similarities with a previously reported mechanism of mammalian PrPSc propagation obtained with a different computational protocol. Together, these results indicate that the propagation of prions generated by evolutionary distant proteins shares common features. In particular, in both these cases, prions propagate their conformation through a very similar templating mechanism.

## Introduction

The phenomenon of protein-based inheritance characterizes prions, proteins appearing at various levels along the evolutionary scale that are capable of propagating their conformationally encoded information in absence of nucleic acids [1]. Despite their original identification as causative agents of neurodegenerative conditions in mammals, prions also exert regulatory roles in different biological contexts [2, 3]. For example, a mechanism of heterokaryon incompatibility in different fungi is regulated by a prion [3–5]. This process reflects the inability of vegetative fungal cells from two different strains to undergo fusion, depending on specific loci (het) whose alleles must be identical for stable hyphal fusion to occur. Strain compatibility ultimately determines whether the heterokaryon develops normally or undergoes controlled cell-death. In *Podospora anserina*, the heterokaryon incompatibility is specified by a het locus appearing as two distinct and incompatible alleles (*HET-s* and *HET-S*), encoding two corresponding proteins (HET-s and HET-S, respectively) [6]. When a HET-s strain fuses with another expressing HET-S, the heterokaryon can undergo controlled cell death. However, incompatibility occurs only when the HET-s factor is folded in an amyloid prion conformation. This trait can be inherited over many generations due to the ability of the HET-s prion to catalyze the conversion of soluble HET-s molecules into the growing amyloid fibrils [7].

The 2-rung-β-solenoid (2RβS) architecture of the HET-s prion has been solved at high resolution by solid-state NMR [8]. On the other hand, the elucidation of the mechanism underlying its propagation is still missing, mainly due to the lack of suitable biophysical methods to characterize such molecular processes at high spatiotemporal resolution.

Computational techniques, such as MD, may in principle be employed to achieve the required level of resolution. However, plain MD simulations can only be applied to study molecular transitions of biologically-relevant proteins occurring in the microsecond timescales [9]. In contrast, folding and misfolding of most proteins and large conformational re-arrangements including prion replication occur at much longer timescales, ranging from milliseconds to minutes [10].

Using enhanced sampling algorithms, it is possible to explore protein conformational states much more efficiently than by plain MD (for recent reviews of some of these methods see e.g. [11, 12]), at the price of having to introduce additional approximations or supply prior information. Yet, most enhanced sampling schemes are still too computationally expensive to be applicable to characterize very rare and complex dynamical processes such as folding, misfolding and amyloidogenic aggregation of most biologically relevant proteins.

An enhanced sampling algorithm that has been successfully applied to investigate the folding and misfolding of several large proteins with realistic all-atom force fields is the so-called Bias Functional (BF) approach [13]. This variational scheme capitalizes on the extremely low computational load of performing rMD simulations [14, 15]. In rMD, an unphysical history-dependent biasing force is introduced to prevent the chain from backtracking along the direction defined by an *arbitrarily chosen* CV. On the other hand, the bias remains latent when the system spontaneously progresses towards the product. The BF variational principle allows the subsequent identification of the rMD transition pathways with the highest probability to occur in the absence of biasing force. It has been rigorously shown that rMD yields realistic transition path ensembles when the CV used in the biasing force is an ideal reaction coordinate (RC), namely the committor function [16]. However, whenever a reasonable proxy of the RC is available, then the BF variational scheme enables to keep the systematic errors of rMD to a minimum. Protein folding pathways obtained with the BF approach have been found to agree very well with the results of both plain MD simulations [13] and kinetic experiments [17, 18], arguably reflecting the fact that a good RC for these is available [19], supported also by energy landscape theory arguments [20]. Unfortunately, the BF approach may be flawed by uncontrolled systematic errors when applied to study processes in which the RC is poorly known.

In our previous work, we used rMD simulations to propose a first full atomistic model for the replication of the mammalian PrPSc prion [21]. To perform such a calculation, first, we obtained an estimate of the reaction coordinate using a phenomenological Markov State Model. Then, we used such a collective variable in the definition of the biasing force. The resulting prion conversion mechanism was one in which PrPC sequentially unfolded and progressively bond to the prion fibril, through a rung-by-rung templated mechanism. An important question to address is to what extent these results depended on our phenomenological estimate of the reaction coordinate. A second important question is whether the templated prion conversion mechanism is universally conserved through different prion species, thus can be extended to HET-s.

In this work, we address these two questions by overcoming the theoretical limitations of our previous study. To this goal, we relied on the much more advanced SCPS algorithm [22]. Like the BF approach, SCPS is powered by rMD-type simulations. However, unlike in BF simulation, the reaction coordinate of SCPS is not heuristically postulated. Instead, it is calculated self-consistently, through an iterative process (S1 Fig). It has been shown that SCPS provides a rigorous mean-field approximation of the unbiased Langevin dynamics, even when the initial guess of reaction coordinate is suboptimal [16, 22]. Thus, SCPS provides a much better tool than BF to investigate structural reactions in which the reaction coordinate is poorly known, thus including prion propagation. The computational cost of SCPS, however, is about one order of magnitude larger.

To date, the SCPS scheme was only tested in a single proof-of-principle simulation of the folding of a 35 amino-acid long protein subdomain, using an implicit solvation model [22]. As a preliminary step, in the Supplementary Information, we report a much more extensive validation, which was performed with an explicit solvent force field on a set 5 different proteins which differ in size, topology and secondary structure composition. Remarkably, the folding pathways of all these chains were found to be statistically indistinguishable from those obtained in the same force field using plain MD simulations on the Anton supercomputer [23].

We then used SCPS scheme to simulate the misfolding of the HET-s prion forming domain and incorporation into a growing fibril. Also in this case, we found that the all-atom reconstruction of the HET-s replication mechanism is characterized by the templated, progressive folding of an unstructured HET-s monomer onto the exposed edges of the growing prion fibrils. Thus, on the qualitative level, these results fully confirm those previously obtained by BF, in our study of mammalian prion replication.

## Results/Discussion

### All-atom reconstruction of HET-s prion propagation

Enhanced path sampling simulations, including SCPS, require a model for the reactant and product states. In our study of HET-s prion propagation, we used as product state the amyloid structure of the HET-s prion forming domain, which includes a trimer previously solved by solid-state NMR (PDB 2KJ3, Fig 1). The monomeric soluble state of the HET-s prion-forming domain is unstructured [24]. Thus, to generate the initial conditions in the reactant state, we performed high-temperature MD simulations on the HET-s trimer introducing positional restraints on heavy atoms on two out of the three monomers. We obtained a total of 10 initial unstructured conditions: 5 initial conditions in which the unstructured monomer resides at the N-terminus (by introducing the restraints on the two C-terminal monomers), and 5 initial conditions in which the unstructured monomer resides at the C-terminus (by introducing the restraints on the two N-terminal monomers; S2 Fig). For each initial condition in the reactant state, we applied the SCPS protocol described in the Methods and retained only the trajectories which successfully reached the product state.

**Fig 1.**
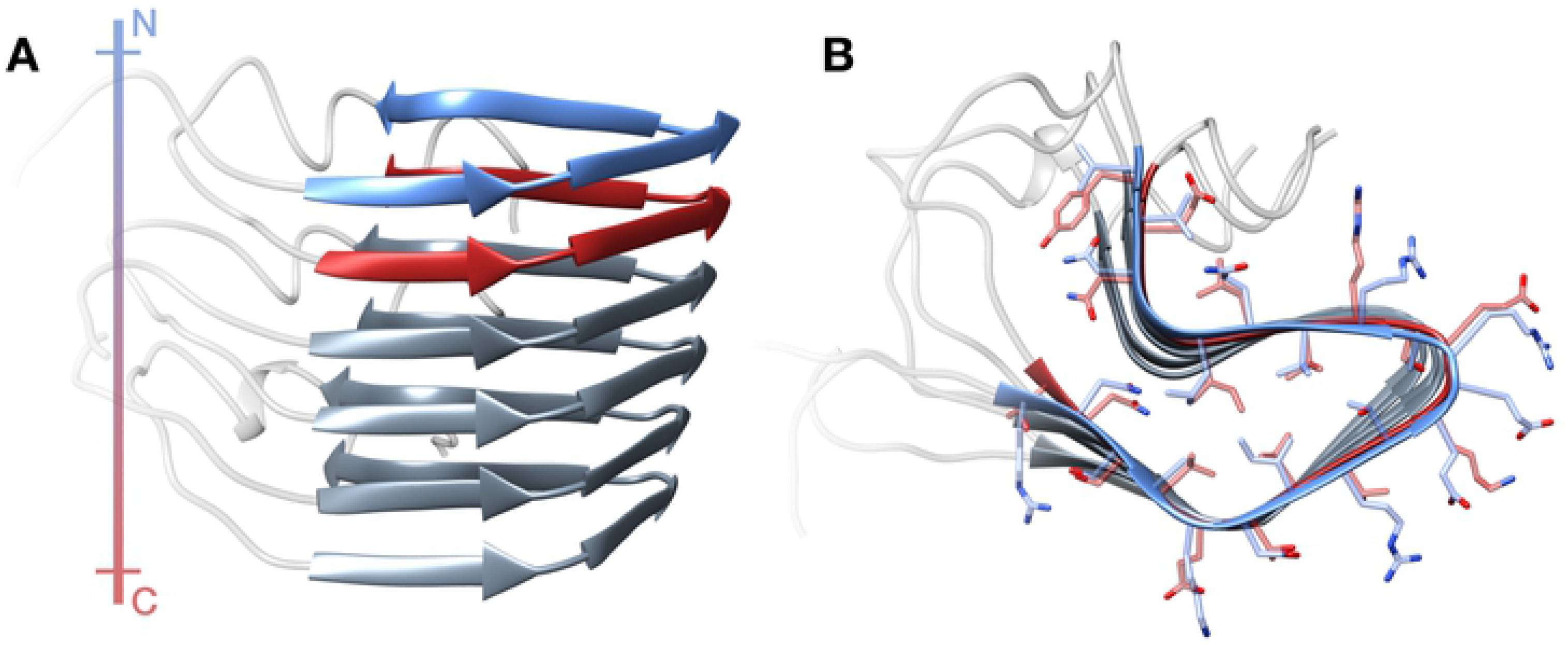
Structure of the HET-s Prion Forming Domain in the amyloid form. Lateral (**A**) and top (**B**) view of a HET-s amyloid trimer retrieved from PDB 2KJ3. Each monomer of the fibril displays a 2RβS conformation. The N-terminal rung (residues 225-245) is depicted in blue, while the C-terminal rung (residues 261-281) is depicted in red. The colored bar at the left represents the polarity (N to C, blue to red) of the fibril.

Let us first discuss the results obtained in rMD simulations, namely before the RC refinement iterations. In the upper- and lower-left panels of Fig 2, we represent the conformations explored by the trajectories projected on the plane defined by the root mean square deviation (RMSD) of atomic positions, to the reference state of each rung of the converting monomer. Namely, the heat map displays the quantity *G(RMSD*_*CT-Rung*_ , *RMSD*_*NT-Rung*_)=−*ln[P(RMSD*_*CT-Rung*_, *RMSD*_*NT-Rung*_*)], where P(RMSD*_*CT-Rung*_, *RMSD*_*NT-Rung*_) is the probability of observing specific pairs RMSD, calculated from a frequency histogram of the rMD trajectories. These results show that the prominent rMD transition path is characterized by the consecutive formation of rungs starting at the fibril ends, as seen from the high-density region near the y-axis. In a few cases, the rungs form simultaneously or with an inverted order.

**Fig 2.**
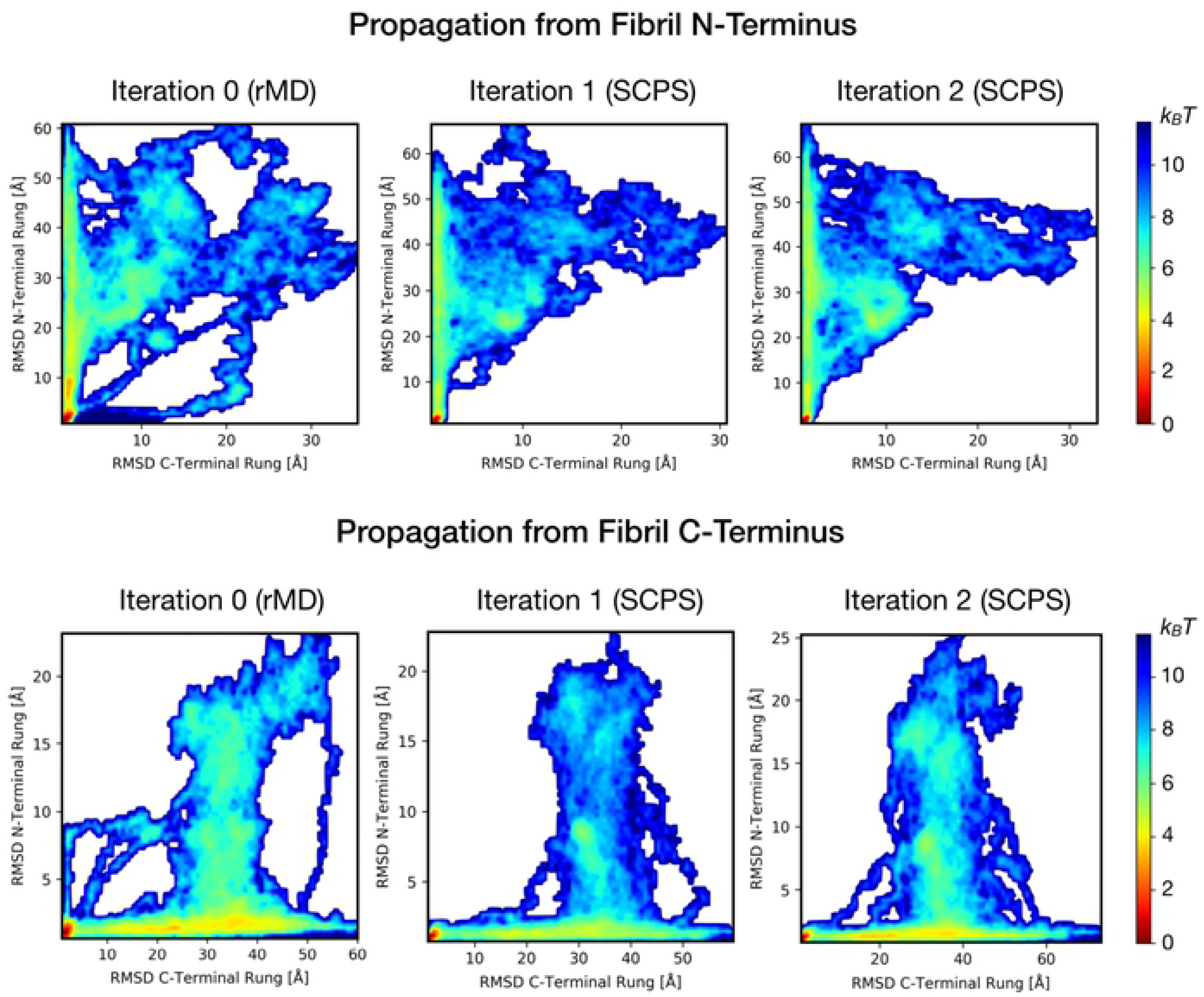
Lower-bound free energy landscapes of HET-s rungs formation. The graphs represent the free energy landscape, computed at each iteration, describing the conformational change of the rungs of the converting HET-s monomer during its incorporation into the fibril. Each landscape was computed as −*ln*[*P*(RMSD_*CT-Rung*_, RMSD_*NT-Rung*_)]. **A**. Graphs relative to trajectories propagating from the fibril N-terminus. **B.** Graphs relative to the trajectories propagating from the fibril C-terminus. In both cases, a prominent pathway, consisting of the consecutive formation of rungs starting at the fibril end, begins to appear in the rMD generated trajectories. However, the rMD algorithm also yields pathways with cooperative or inverted rung formation. These alternative events are completely absent after a single SCPS iteration. Of note, a second SCPS iteration does not produce a consistent change in the free energy landscape in both A and B, indicating the convergence of the algorithm.

After a single SCPS iteration, the entire set of productive trajectories converged on a single reaction path, which proceeds by forming the rung anchoring the fibril end, followed by the formation of the subsequent rung. A second SCPS iteration did not produce significant alterations of the energy landscapes, indicating the convergence of the algorithm. The disappearance of secondary propagation pathways after employing the SCPS suggests that their sampling was the result of a non-ideal initial reaction coordinate. These data support a picture according to which the process of HET-s prion propagation occurs by a templating mechanism (S1. Movie and S2 Movie).

To further characterize the reaction process underlying the templated conversion of HET-s prions, we performed additional analyses. In Fig 3, we show the median value of the reaction progress variable *Q* which each residue assumes the β-strand conformation. Residues forming at high values of *Q* assume β-strand conformation in the late stage of the reaction. This analysis shows that, in the case of initial anchoring occurring at the N-terminus of the HET-s fibril, the C-terminus of the incoming monomer acquires a β-sheet conformation by progressively establishing intermolecular hydrogen bonds with the exposed edge residues. Once completed, the new rung templates the formation of intramolecular hydrogen bonds with the remaining residues of the polypeptide. Insertion of monomers at the C-terminal edge of the HET-s fibril proceeds in a specular fashion. These observations indicate that the two exposed edges of the HET-s amyloid provide the initial scaffold for the templated conversion of incoming monomers, which become new edges after completion of the reaction.

**Fig 3.**
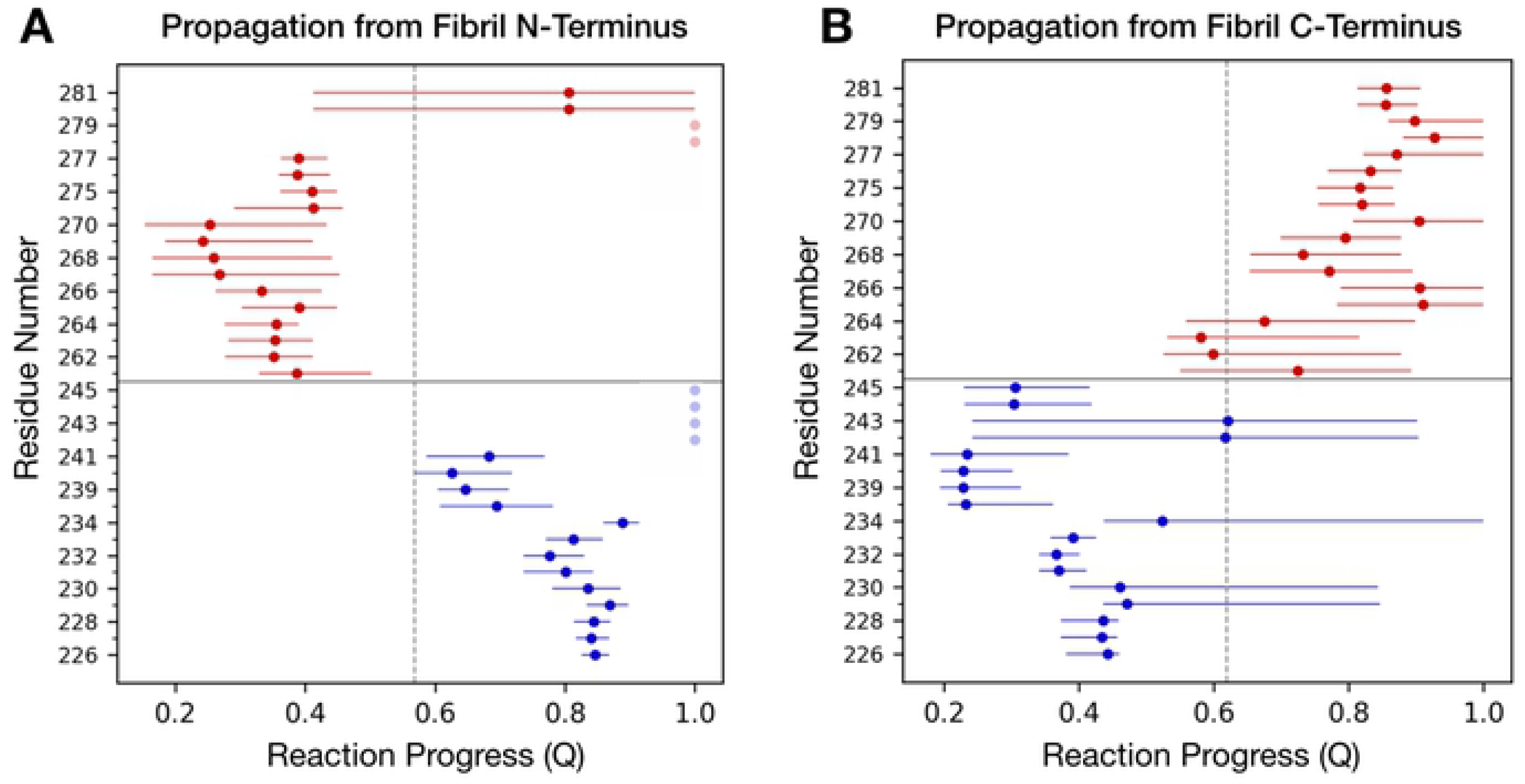
Order of β-strand formation along the HET-s propagation pathway. The plots show the median reaction progress at which each residue of the rungs assumes the β-strand conformation. Blue dots correspond to residues in the N-terminal rung while red dots correspond to residues in the C-terminal rung. Dots shown in transparency indicate residues not achieving a stable β-strand conformation. Horizontal bars span between the first and the third quartile of the distribution. The vertical dashed line delineates the average reaction progress at which half of the rungs-residues are incorporated into β-strand conformation. In the propagation starting from the fibril N-terminus (**A**) the residues of the C-terminal rung of the converting monomer are incorporated first (mean Q = 0.44), followed by the residues in the N-terminal rung (mean Q = 0.69). On the contrary, in the propagation starting from the fibril C-terminus (**B**) the residues of the N-terminal rung of the converting monomer are incorporated first (mean Q = 0.42), followed by the residues in the C-terminal rung (mean Q = 0.81).

To obtain an unbiased projection of the reaction paths without selecting *a priori* the collective variables for its representation, we performed a principal component analysis (PCA) on the Cα contact maps of the sampled transition pathways (Fig 4A and 4D). The PCA energy landscape representation showed that in both N-terminal and C-terminal propagation pathways the reaction initially occurs almost exclusively along the principal component 2 (PC2), and only subsequently toward the principal component 1 (PC1). Further analysis of the contribution of each contact distance to the two principal components was performed by grouping the contacts in three sets: (i) contacts between residues belonging to the same rung (intra-rung); (ii) contacts between residues of different rungs (inter-rung); (iii) contacts between residues of the converting monomer and the structured fibril (monomer-fibril) (Fig 4B and 4D). This analysis revealed that the main contact contributors for PC1 belong to the inter-rung set, while contributors of PC2 belong mainly to the monomer-fibril set. The PCA analysis repeated using an all-atom contact map yielded to overlapping results (S3 Fig).

**Fig 4.**
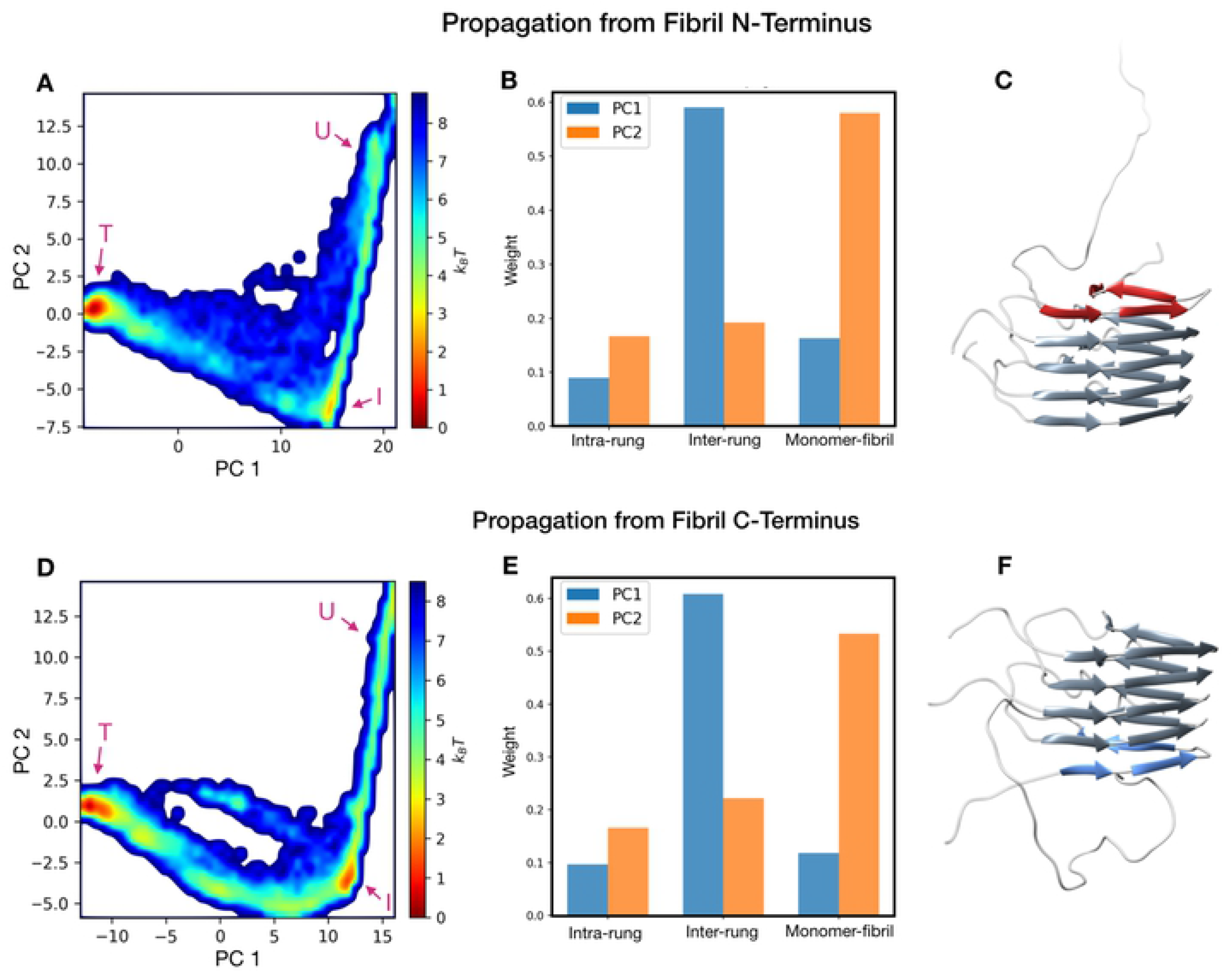
Principal component analysis of the HET-s propagation trajectories. Graphs (**A**) and (**D**) represent the free energy landscape in the principal component plane of the trajectories propagating from the fibril N- and C-terminus respectively. The letters “U”, “I” and “T” indicate the unstructured, intermediate the final (target) state, respectively. The residue contacts were classified in three categories: contacts between Cα in the same rung (intra-rung), contacts between Cα belonging to different rungs (inter-rung) and contacts between Cα of the converting monomer and Cα of the structured fibril (monomer-fibril). The contribution of these sets of the two principal components is shown in bar plots (**B**) for the N-terminal propagation and (**D**) for the C-terminal propagation. Images (**C**) and (**F**) show representative protein conformations sampled from the intermediate state, for the N-terminal and C-terminal propagations, respectively.

Overall, these findings corroborate a rung-by-rung propagation model for HET-s. Interestingly a highly populated region appears at the elbow of the landscape graph, in both the N-terminal and C-terminal fibril growths. Such a region reflects long-living misfolding intermediates occurring during the HET-s templated conversion mechanism, characterized by the presence of only one rung of the converting monomer attached to the elongating fibril (Fig 4C and 4F).

Collectively, our SCPS simulations indicate that HET-s prion conversion occurs through a series of templating events occurring at the N-terminal and C-terminal ends of the fibrils. In these sites, incoming HET-s monomers are initially incorporated by establishing intermolecular hydrogen bonds with the exposed β-strands, leading to the formation of a first rung, which then forms new intramolecular hydrogen bonds with the remaining parts of the polypeptide, giving rise to a new 2RβS fibril subunit (Fig 5).

**Fig 5.**
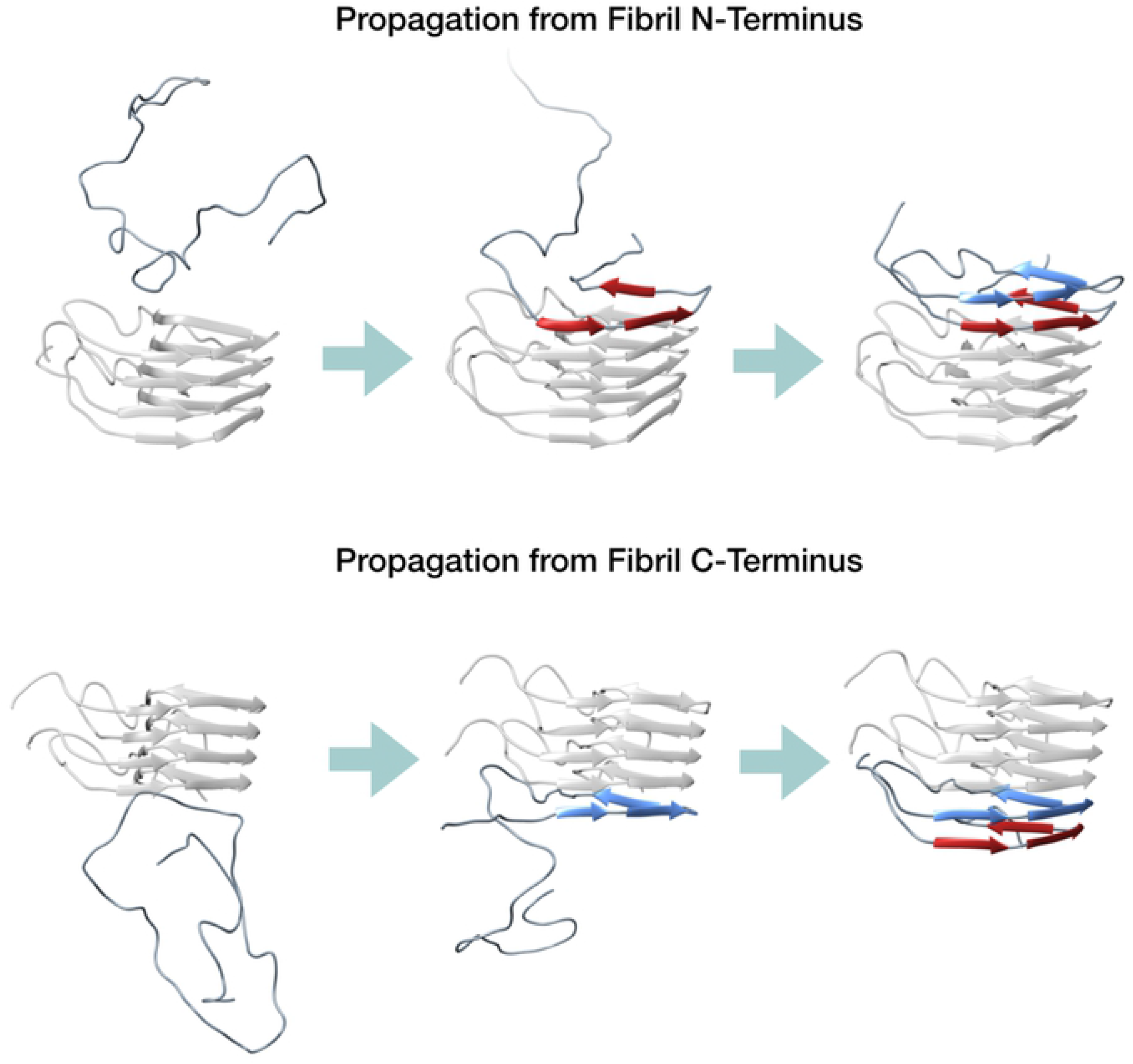
Representative scheme for HET-s prion propagation. **A.**Propagation scheme starting from the N-terminal end of the fibril. In this process, the C-terminal rung of the converting monomer (depicted in red) is formed by templating onto the structured N-terminal rung of the fibril (shown in transparent grey). Subsequently, the N-terminal part of the converting monomer forms the second rung (depicted in blue); **B**. Propagation scheme starting from the C-terminal end of the fibril. In this process, the N-terminal rung of the converting monomer (depicted in blue) is formed by templating onto the structured C-terminal rung of the fibril (shown in transparent grey). Subsequently, the C-terminal part of the converting monomer forms the second rung (depicted in red).

We emphasize that the results of our SCPS simulation of the entire sequence of events underlying the incorporation of HET-s monomers into fibrils were obtained without relying on a specific choice of RC, i.e. the absence of strong ad-hoc assumptions concerning the reaction mechanism.

From a statistical mechanics perspective, the results of our numerical simulations can be rationalized by noting that the presence of a seeding template strongly reduces the free-energy barrier for a misfolding transition, thus effectively activates the fibrilization reaction [25]. These numerical and theoretical results are in line with experimental data, which show that in seeds-free conditions, the spontaneous fibrillization of soluble HET-s conformers is characterized by a lag phase (nucleation process), which is instead abrogated in the presence of HET-s prion seeds [26]. Consistent with our propagation model, the seeds can provide the templating surfaces that effectively catalyze the incorporation of new monomers at the fibril ends.

### Comparison between the replication mechanism of PrP^Sc^ and HET-s prions

The main difference between the replication mechanisms of PrP^Sc^ and HET-s prion in our simulations is the monodirectional growth of the former in contrast to the bidirectional extension of the latter. The reason for such difference is related to the conformation of the monomers acting as substrates. The propagation of PrP^Sc^ involves the refolding of both structured (C-terminus) and unstructured (N-terminus) domains of PrP^C^, while the propagation of HET-s involves exclusively the intrinsically disordered prion-forming domain of the protein. Such a difference implies that, while the replication mechanism of PrP^Sc^ has an intrinsic directionality determined by the initial binding and subsequent refolding of parts of the unstructured N-terminal domain, the HET-s prion may replicate by engaging interactions with both the N-terminus or C-terminus of its substrate. Regardless of such a difference, PrP^Sc^ and HET-s prions share an almost identical propagation mechanism, characterized by a progressive, rung-by-rung pathway, governed by the cooperativity of hydrogen bonds and the lateral stacking of the side chains.

Overall, the results reported in this work show that refined computational tools like the SCPS can provide valuable insight into the atomistic details of misfolding events, that may inspire original hypotheses laying the groundwork for new experimental approaches. The concordance between the results of our computational studies on HET-s and PrP^Sc^ provides the first evidence that prions from distant species may propagate similarly, implying that this common prion replication mechanism could have been conserved through evolution.

## Methods

### Implementation of Self-Consistent Path Sampling

SCPS is an iterative enhanced path sampling algorithm derived directly from the Langevin equation which can be used to effectively generate ensembles of transition pathways connecting a given ensemble of configurations to configurations in a target state. Its implementation consists of three steps (S1 Fig).

#### STEP 1: rMD simulations

Starting from a given initial condition in the reactant, an ensemble of reactive trajectories is generated using the rMD algorithm. In these simulations, an external biasing force is introduced to impair backtracking towards previously visited states defined along a single reaction coordinate *z(X)*. The biasing force acting on a single atom is computed as:

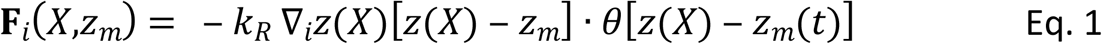

where *k*_*R*_ determines the strength of the biasing force and *z(X)* is the CV which represents an initial guess for the reaction coordinate. In the specific case of protein folding simulations and the present simulation of HET-s propagation, we adopted a definition of *z(X)* based on the overlap between the instantaneous contact map and the target contact map:

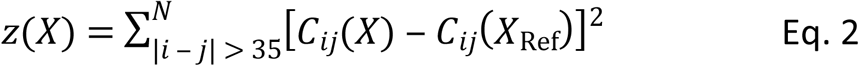

In this equation, X_Ref_, is a reference three-dimensional structure in the target state and the constraint |i-j|> 35 is introduced to exclude the trivial contacts due to bonded interactions within amino-acids. The entries of the contact map are a continuous function of atomic configuration, defined as:

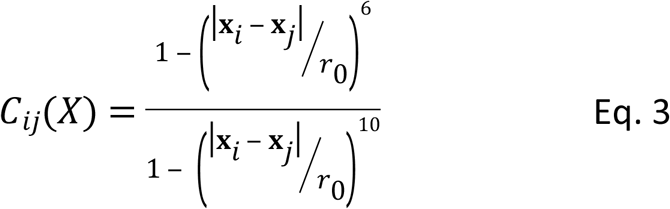

where *r*_*0*_ is a reference distance, set to 7.5 Å, while **x**_*i*_ and **x**_*j*_ are the atomic coordinates of atoms *i* and *j*, respectively. In equation 1, the term *z*_*m*_*(t)* indicates the minimum value assumed by *z(X)* up to the previous integration time-step, i.e. the maximum overlap with the target structure. Therefore, the external force is introduced only when the system backtracks toward the reactant state (i.e. when *z(X)* > *z*_*m*_), otherwise, the system spontaneously proceeds as in plain MD. At the end of the simulation, a trajectory is considered to have successfully reached the target state if a frame *n* satisfies the following conditions:

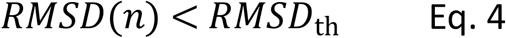

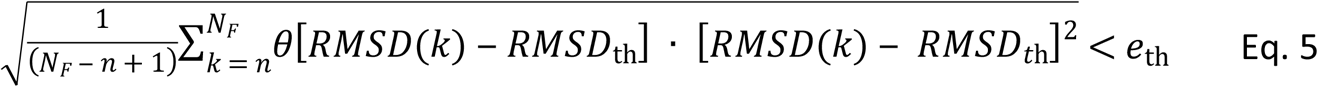

The RMSD to the target reference state is computed for each frame and *RMSD*_th_ represents the closeness threshold, which was set to 3 Å. In equation 5, *N*_*F*_ is the total number of trajectory frames and *e*_th_ is a tolerance threshold. Therefore, to properly consider a trajectory reaching the target reference state, one of its frames must reach below the closeness threshold. The root-mean-squared error, computed on the subsequent frames that are above the threshold, must lay below the tolerance *e*_th_, that was set to 0.3 Å.

#### STEP 2: Calculation of the Mean Transition Paths

The contact maps entries *C*_*ij*_*(X)* calculated frames of all the rMD trajectories reaching the target state starting from an identical initial conformation are averaged to estimate their iso-time mean, resulting in the mean-path of the folding process:

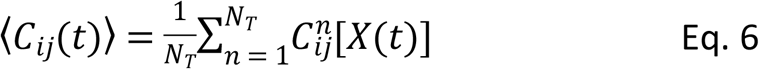

Here *C*_*ij*_^*n*^*[X(t)]* is an element of the contact map computed at time *t* for the *n*_*th*_ trajectory, and *N*_*T*_ is the total number of successful reactive trajectories generated from that initial condition. For convenience, the mean path is downsampled to a low number of contact maps equally spaced in *z(X)* distance, namely ⟨*C* ⟩_*k=1 … NC*_.

The mean path is then used to define two new coordinates:

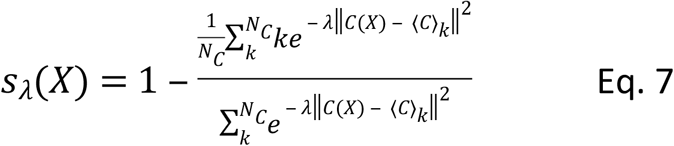

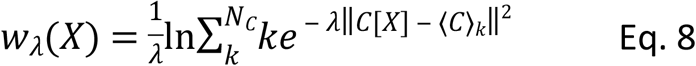

where ||…|| denotes the norm defined in Eq. 2, *⟨C⟩*_*k*_ is the *k*_*th*_ contact map along the mean path, *N*_*C*_ is the total number of contact maps and *λ* is a parameter which is usually set to the inverse of the average distance between those contact maps. The former collective variable represents the progress of the reaction taking the mean path as reference. By definition *s*_*λ*_*(t)* is 1 in the unfolded state and 0 in the native state.

The second coordinate *w*_*λ*_*(X)* measures the shortest distance of the configuration *X* from points in the mean path.

#### STEP 3: Self-Consistent Refinement of the RC

A new set of folding trajectories is generated by employing a modified version of the rMD algorithm, introducing two biasing forces analog to the one defined in Eq. 1, but using *s*_*λ*_*(X)*, and *w*_*λ*_*(X)* instead of *z(X)* as collective variables, respectively. The productive trajectories generated this way (as defined in Eq. 4 and 5) are then used to compute a new mean path. In turn, an updated version of the *s*_*λ*_*(X)*, and *w*_*λ*_*(X)* collective variables.

Steps 2 and 3 are iterated until convergence, that is when a new iteration produces identical results to the previous one, according to some arbitrary convergence criterion. In this work, we adopted one based on the structure of reaction pathways in the plane selected by the RMSD to the native structure of the two terminal rungs (see Fig. 2).

### Simulations of HET-s propagation

SCPS simulations of HET-s propagation were performed using the Charmm36m force field and the modified Charmm TIP3P water model [27]. Initial conformations, each one consisting of a HET-s dimer in the amyloid form and an unstructured monomer, were generated by thermal denaturation. In particular, the trimeric amyloid structure (PDB 2KJ3) was positioned in a cubic box with an approximate side length greater than 120 Å that was filled with TIP3P water molecules. The system was neutralized with 3 Cl^−^ ions and energy minimized with the steepest descent method. Then, 200 ps of NVT equilibration using the V-rescale thermostat at 800 K were carried out by employing position restraints with a force constant of 103 kJ·mol^−1^·nm^−2^ on heavy atoms. Finally, 5 simulations were performed for 2 ns each at 800 K by releasing the restraints on the N-terminal monomer and the other 5 by releasing the restraints on the C-terminal monomer (S2 Fig.). The sampled initial conformations were re-solvated in a smaller cubic box with an approximate side of 100 Å; then 3 Cl^−^ ions were added to counterbalance the protein charge and the system was brought to a final 150 mM NaCl concentration. An energy minimization, 500 ps of NVT (310 K) and 500 ps of NPT (310 K, 1 Bar) equilibrations were carried out for each condition with position restraints on heavy atoms. The V-rescale thermostat and the Parrinello-Rahman barostat were employed for equilibrations and the subsequent rMD and SCPS production. First, 20 rMD simulations for each initial condition were performed, consisting of 3 · 10^6^ steps employing the leap-frog integrator with 2 fs time-step. Frames were saved every 500 steps. The value of *k*_*R*_ was set to 5 · 10−5 kJ mol^−1^ and *r*_*0*_ to 7.5 Å. The cutoff radius to ignore contact pairs was 12 Å. The trajectories successfully reaching the target state according to Eq. 4 and 5 were used to compute the mean path of each condition (Eq. 6), which was down-sampled to 10 equally spaced contact maps. From each initial condition, 20 simulations were then re-launched for SCPS with bias constants *k*_*s*_ = 1.5 10^−5^ kJ mol^−1^ and *k*_*w*_ = 3 · 10^−5^ kJ mol^−1^. A total of 2 SCPS iterations for each initial condition were performed.

### Analysis of HET-s simulations

The analysis regarding the order of rung formation was first performed by computing the RMSD to the target state of the rungs of the converting monomer. The all-atom RMSD of the two rungs (N-term: R225 to V245, C-term: T261 to Y281, rungs depicted in Fig. 1) was evaluated for all the trajectories successfully reaching the target state. The calculation of the RMSD was performed with alignment on the two monomers in the amyloid state (the converting monomer was not included for the alignment). For each iteration, two different energy landscapes were generated using frequency histograms: the first landscape was constructed using all the productive trajectories in which the HET-s propagation starts at the N-terminus of the fibril. The second landscape was obtained collecting all the productive paths in which the HET-s propagation starts at the C-terminus of the fibril. It is important to emphasize that these landscapes are not directly related to free energy distribution, because our reactive trajectories do not sample the Boltzmann distribution. Instead, these frequency histograms measure the probability of visiting a given configuration along the calculated transition paths. The median point in the reaction progress at which each residue of the converting monomer assumes a β-strand structure was computed. A residue was considered to form a β-strand the first time that such conformation was kept for more than 5 frames consecutively. The time of formation was then converted to the corresponding value of Q (corresponding to the fraction of reference contacts). Calculations of the secondary structures were performed with the STRIDE algorithm. Finally, PCA analysis was performed by using the Cα contact maps and the all-atom contact maps (excluding hydrogens). For the N-terminal propagation simulations, the contact maps were calculated using all the residues of the converting monomer and the residues belonging to the N-terminal rung of the N-terminal monomer already included in the fibril in the initial state. For the C-terminal propagation simulations, residues of the converting monomer and the C-terminal rung of the C-terminal monomer already included in the fibril were used for contact map calculation. In this all-atom PCA analysis, each trajectory was downsampled to a total of 300 frames.

### Data production, analysis, and visualization software

Biased simulations (rMD and SCPS) were performed in Gromacs 2018 [28], where we implemented the collective variables *z(X)*, *s*_*λ*_*(t)* and *w*_*λ*_*(t)*. Data analysis was performed in python using the following libraries: MDAnalysis, NumPy and SciPy. Python scripts were accelerated with the Numba compiler. Graphs were obtained by using Matplotlib in python. Images of protein conformations were generated in UCSF Chimera.

## Supplementary Figure Legends

**S1 Fig. Schematic representation of the SCPS algorithm.**

In this figure, *X*_*I*_ represents the initial protein conformation, *X*_*F*_ the final target conformation and T_1…N_ the trajectories connecting *X*_*I*_ to *X*_*F*_. **Step 1:** an ensemble of trajectories starting from *X*_*I*_ and successfully reaching *X*_*F*_ is generated by employing the rMD, biasing along the pre-defined *z(X)* reaction coordinate. Since *z(X)* decreases when the proximity of *X* to *X*_*F*_ increases, the progression along the path is here defined as *−z(X)*. **Step 2:** the trajectories successfully reaching the target state are then used to compute the mean path *⟨C*_*ij*_*(t)⟩*, depicted in purple. The mean path is then used to define two new coordinates: *s*_*λ*_, depicted in blue, which value is 1 in the unstructured state and 0 in the target state, therefore (1 - *s*_*λ*_) is used here to define the progress along the mean path; and *w*_*λ*_, depicted in red, that represents the distance to the mean path. **Step 3:** a modified version of the rMD is employed to generate a new set of trajectories by introducing two biasing forces, acting along *s*_*λ*_*(t)*, and *w*_*λ*_*(t)* instead of *z(X)*. The trajectories successfully reaching the target state are then used to compute a new mean path (step 2) to perform a new iteration.

**S2 Fig. Initial conditions of HET-s simulations.**

The initial conditions used to generate the propagation pathways are reported. Initial conditions for simulating propagation from the fibril N-terminus were obtained by performing high-temperature MD, introducing positional restraints on heavy atoms on the two C-terminal monomers. Initial conditions for simulating propagation from the fibril C-terminus were obtained by performing high-temperature MD, introducing positional restraints on heavy atoms on the two N-terminal monomers.

**S3 Fig. PCA of the HET-s propagation trajectories using all no-H atoms.**

Graphs on the left represent the free energy landscape in the principal component plane of the trajectories propagating from the fibril N-terminus (**A**) and C-terminus (**B**), respectively. Bar plots on the right show the contribution of the contact-type sets for the two principal components.

## Supplementary Movie Legends

**S1 Movie. Propagation of the Het-s prion from the N-terminal surface of the fibril.**

Visualization of a representative SCPS trajectory of a Het-s monomer converting at the N-terminal surface of the fibril.

**S1 Movie. Propagation of the Het-s prion from the C-terminal surface of the fibril.**

Visualization of a representative SCPS trajectory of a Het-s monomer converting at the C-terminal surface of the fibril.

## Acknowledgements

This work was supported by a grant from Fondazione Telethon (Italy, TCP14009). GS is a recipient of a fellowship from Fondazione Telethon. EB is an Assistant Telethon Scientist at the Dulbecco Telethon Institute (Fondazione Telethon, Italy).

## Author Contribution

Conceptualization: GS, JRR, EB and PF

Data curation: LT, GS and AB

Formal analysis: LT, GS, AB and PF

Funding Acquisition: EB and PF

Investigation: LT, GS, AB, JRR, EB and PF

Methodology: LT, GS, AB, EB and PF

Project Administration: EB and PF

Software: LT, GS and AB

Supervision: EB and PF

Visualization: LT and GS

Writing, original draft: GS, EB and PF

Writing, review and editing: LT, AB and JRR

## Competing Interests

LT and AB have direct involvement in the ongoing research at Sibylla Biotech SRL (www.sibyllabiotech.it). GS, EB and PF are co-founders and shareholders of the company.

